# Assessment of the need for separate test set and number of medical images necessary for deep learning: a sub-sampling study

**DOI:** 10.1101/196659

**Authors:** Ariel Rokem, Yue Wu, Aaron Lee

**Affiliations:** The eScience Institute, University of Washington; University of Washington, Department of Ophthalmology

**Keywords:** Opthalmology, Retina, Optical Coherence Tomography, Macula, Deep learning, Machine Learning

## Abstract

Deep learning algorithms have tremendous potential utility in the classification of biomedical images. For example, images acquired with retinal optical coherence tomography (OCT) can be used to accurately classify patients with adult macular degeneration (AMD), and distinguish them from healthy control patients. However, previous research has suggested that large amounts of data are required in order to train deep learning algorithms, because of the large number of parameters that need to be fit. Here, we show that a moderate amount of data (data from approximately 1,800 patients) may be enough to reach close-to-maximal performance in the classification of AMD patients from OCT images. These results suggest that deep learning algorithms can be trained on moderate amounts of data, provided that images are relatively homogenous, and the effective number of parameters is sufficiently small. Furthermore, we demonstrate that in this application, cross-validation with a separate test set that is not used in any part of the training does not differ substantially from cross-validation with a validation data-set used to determine the optimal stopping point for training.

## I. Introduction

DEEP learning (DL) algorithms [1] have been tremendously successful at solving a variety of different computational tasks. Although these algorithms were originally developed to perform computer vision tasks that require the identification and classification of natural objects in images [2], they have been more recently successfully applied in tasks as varied as automated captioning and description of images and videos [3], [4], automated transcription of spoken language [5] automated translation [6], and in playing games such as Go [7] and Poker [8], beating even highly seasoned players in these games.

A key to the success of these algorithms in image processing is that they do not require feature engineering of their front-end filters. Instead, these filters are empirically learned from the data through a process of training. For example, a network is trained to identify natural objects by exposing it to labeled exemplars of images containing the classes to be discriminated. The weights that define the front-end filters, and their pooling in higher levels are automatically adjusted through gradient descent. Progress in the implementation of the computation of the gradients required for this process on graphical processing units (GPUs) has been instrumental in enabling use of these techniques on large and complex data-sets. No less important have been the discovery of network architectures [9], nonlinearities [10], regularization procedures [11] and initialization procedures [12] that accelerate and improve learning. Taken together, these factors have ushered in an era where training of large DL networks has become practical, and there is wide-spread interest in applying these algorithms to a variety of different tasks.

### A. Deep Learning in medical imaging

Because DL algorithms were originally developed to perform difficult image processing and classification tasks, one of the compelling avenues for application of DL is in the analysis of data from medical imaging technologies, and the development of computer-assisted diagnostic systems with DL-trained networks at their core. This type of application is rapidly becoming more realistic because of the combination of high-quality biomedical imaging technologies that are becoming common in clinical practice, and the development of large data-sets for the training of these networks. These large data-sets are the result of years of accumulation of electronic medical records (EMR), data-bases that include both image data, as well as expert-generated diagnostic labels. Several recent studies used data from such data-bases in tandem with DL networks to demonstrate the potential for highly accurate automated diagnosis of diseases from images of retina [13] and of skin [14].

In previous work, we demonstrated that a DL algorithm is capable of accurate diagnosis of age-related macular degeneration (AMD) from images of the retina acquired with optical coherence tomography (OCT) [15]. OCT images are taken at a high resolution, and provide information about the three-dimensional structure of the tissue. They routinely collected during clinical ophthalmological visits, and are normally used by clinicians to assess the presence of retinal diseases. They are stored in an EMR, and the combination of the diagnostic labels available in the EMR database, together with the images of retinae from both healthy individuals and AMD patients allowed us to train a DL network with the VGG16 architecture, previously used for object recognition [16], to distinguish OCT scans from retinae of patients with AMD from healthy retinae with an accuracy of 88.98% (ROC AUC of 93.83%, peak sensitivity 92.64 %, peak specificity 93.69 %).

### B. How many samples do you need?

Previous applications of DL in medical imaging, including our own work, relied on large data sets, that are not available for many other technologies, and for diseases that are less common. A major barrier to the wide-spread application of DL algorithms in medical imaging is the assumption that these algorithms only work well when data is extremely abundant, and that supervised learning can progress using accurate labels of each image^1^. Indeed, in our previous work using DL for OCT image classification, the network was trained with a data-set of ~100,000 images. Similarly, other studies have used data-bases with many thousands of patients and up to millions of individual images.

To our knowledge, there is only one previous study asking how many samples are needed in biomedical image classification [18]. The authors of this study trained a DL network to discriminate between six classes of images (brain, neck, shoulder, etc.) from MRI images. They found that only a few hundred images are required to reach near-perfect accuracy in this task using the GoogLeNet network. However, images of these body parts differ in many respects, and it is not clear that a much simpler algorithm would perform just as well in this classification task.

In the present work we focused on a classification task in which DL algorithms are required to perform more accurately than traditional image processing methods [19]. We introduce a resampling procedure to test the size of the sample needed in order to train a DL network on a biomedical image classification task, and use this procedure in order to assess the number of samples needed to train a network to accurately discriminate between AMD and healthy retinae from OCT.

### C. Cross-validation and the importance of a separate test set

To avoid over-fitting, and to provide an objective and accurate evaluation of the performance of a classification algorithm, it is common to separate the data into several different sets: a *training set* is used to learn the dependencies between input data and model class labels, and to adjust the parameters of the model. A *validation set* is sometimes used to assess the current state of the model during training. This is done by feeding a sample or samples from the validation set through the algorithm, with a fixed set of parameters, and evaluating the accuracy of the classification with these parameter values, but without using the results to adjust the parameters.

Often, an additional data set is set aside as *test set*. This set is used once the learning has ended, as a single independent estimate of the endpoint of learning. While using an independent data set completely guards against the danger of over-fitting, it also might introduce the danger of variability in the estimate of error, especially with a relatively small size of the test set [20].

In cross-validation, different parts of the data might serve separately as training and validation data sets [21]. For example, in k-fold cross-validation training is repeated several times, where in each iteration through the procedure, a portion of *N/k* samples from the data are designated as a validation set, and the remaining data is used for training. After *k* repetitions of this procedure, all the data has been used up as validation data. This means that a full set of errors on the entire data-set has been computed. This procedure is sometimes used for comparative evaluation of different models, and for model selection (by comparing cross-validation errors for two or more models) [22].

While this procedure is comprehensive, and potentially reduces variability of the estimates, it is also computationally demanding. For this reason, training of DL algorithms usually uses the strategy of training on a sub-set of the data and then using other sub-sets for evaluation and testing. Furthermore, given the large amounts of data that are often available for training and evaluation, the final test step is often omitted in applications of DL. For example, in their highly influential paper on image classification from the ImageNet dataset, Krizhevsky et al. [23] comment that: “ In the remainder of this paragraph, we use validation and test error rates interchangeably because in our experience they do not differ by more than 0.1%”.

The resampling scheme introduced here also presents an opportunity to evaluate whether this statement generalizes well to situations in which data is much less abundant. Therefore, a second aim of the present paper is to assess the use of a separate test in cross-validation of a DL network.

## II. Methods

This study was approved by the Institutional Review Board of the University of Washington (UW) and adhered to the tenets of the Declaration of Helsinki and the Health Insurance Portability and Accountability Act.

### A. Optical Coherence Tomography Imaging and Electronic Medical Record Extraction

Macular OCT scans were acquired in the course of clinical care, using a Heidelberg Spectralis OCT scanner (Heidelberg Engineering, Heidelberg, Germany). High-resolution images of the retinal cross-section were obtained using a 61-line raster scan. All of the images from the period 2006 to 2016 were extracted using an automated extraction tool from the instrument imaging database. The images were linked by patient medical record number and dates to the clinical data stored in EPIC. Specifically, all clinical diagnoses and the dates of every clinical encounter, macular laser procedure, and intravitreal injection were extracted from the EPIC Clarity tables.

### B. Patient and Image Selection

A normal patient was defined as having no retinal International Classification of Diseases, 9th Revision (ICD-9) diagnosis and better than 20/30 vision in both eyes during entirety of their recorded clinical history at UW. An AMD patient was defined as having an ICD-9 diagnosis of AMD (codes 362.50, 362.51, and 362.52) by a retina specialist, at least 1 intravitreal injection in either eye, and worse than 20/30 vision in the better-seeing eye. Patients with other macular pathology by ICD-9 code were excluded. These parameters were chosen *a priori* to ensure that macular pathology was most likely present in both eyes in the AMD patients and absent in both eyes in the normal patients. Consecutive images of patients meeting these criteria were included, and no images were excluded due to image quality. Labels from the EMR were then linked to the OCT macular images, and the data were stripped of all protected health identifiers.

As most of the macular pathology is concentrated in the foveal region, the decision was made *a priori* to select the central 11 images from each macular OCT set, and each image was then treated independently, and labeled as either normal or AMD. The images were histogram equalized and the resolution down-sampled to 192 by 124 pixels to accommodate RAM limitations.

### C. Deep Learning Classification Model

A modified version of the VGG16 convolutional neural network [16] was implemented using Caffe [24]. This network was originally designed to classify categories in natural images and was adapted here to classify healthy and AMD retinae. Weights were initialized using the Xavier algorithm [12]. Training was then performed using multiple iterations, each with a batch size of 100 images. ADAM optimization [25] was used with a starting learning rate of 2 × 10^-7^ and the momentum parameters set to 0.9 and 0.99. The loss of the model was recorded at each training iteration, and cross-validation with a separate validation set was conducted every 250 iterations. The training was stopped when the loss of the model decreased and the accuracy of the validation set also decreased (indicating that the model was in the over-fitting regime).

### D. Sub-sampling experiments

At the outset of the experiments, a random subset of 10% of the images were segregated at the patient level into a separate *test set* of images. These would be used to test the performance of the DL network at the end of training. The remaining 90% of the images were then segregated into 11 replicates of random subsets of 4%, 8%, 16%, 32%, 64%, and 100% of the available images. Within each subset, the images were again subdivided into 75% for training and 25% for validation. Care was taken to ensure that the validation set and the training set contained images from a mutually exclusive group of patients (ie, no single patient contributed images to both the training and validation sets). The order of images of the training set was then randomized in each replication condition.

Each replication condition was then trained for a total of 75,000 iterations and the maximal validation accuracy was recorded. The weights at the time of the maximal validation accuracy was used to assess the performance of the network against the held-out test set.

## III. Results

### A. Data and subsampling

More than 2.6 million optical coherence tomography (OCT) images were extracted from the imaging database and linked to clinical data from the electronica medical records (EMR). A total of 48,312 normal OCT scans and 52,690 AMD scans met the inclusion criteria for use in the training set. At the outset, a test set was set aside comprising of 9,493 images, to be used only for evaluation of the training procedure once it is done.

To test the effect of sample size on accuracy of classification with a DL network, random subsets of 4%, 8%, 16%, 32%, 64%, and 100% were created from the full dataset, and training was conducted using these different subset sizes. The breakdown in the number of images in each subset is described in Table I.

**TABLE I.**
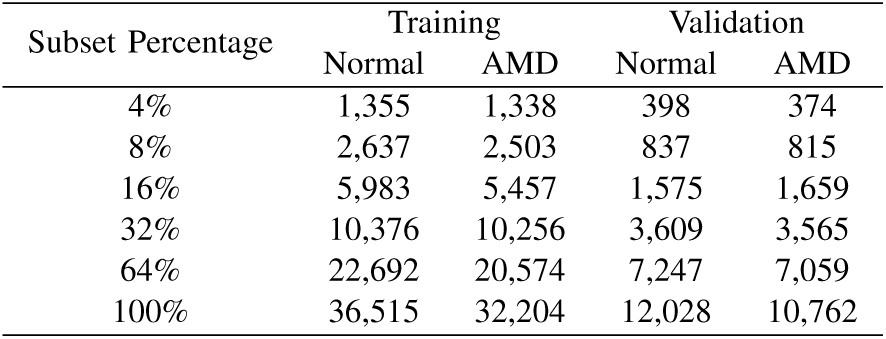
Average Image Counts for Each Subset Condition

### B. Learning with different size subsamples

To assess the robustness of the results to the random selection of specific images, random subsamples of each one of these proportions were drawn from the full data-set 11 times. Training of the DL network in each repetition was allowed to progress for 75,000 iterations. Learning curves, recording validation accuracy during training are shown in Figure 1.

**Fig. 1.**
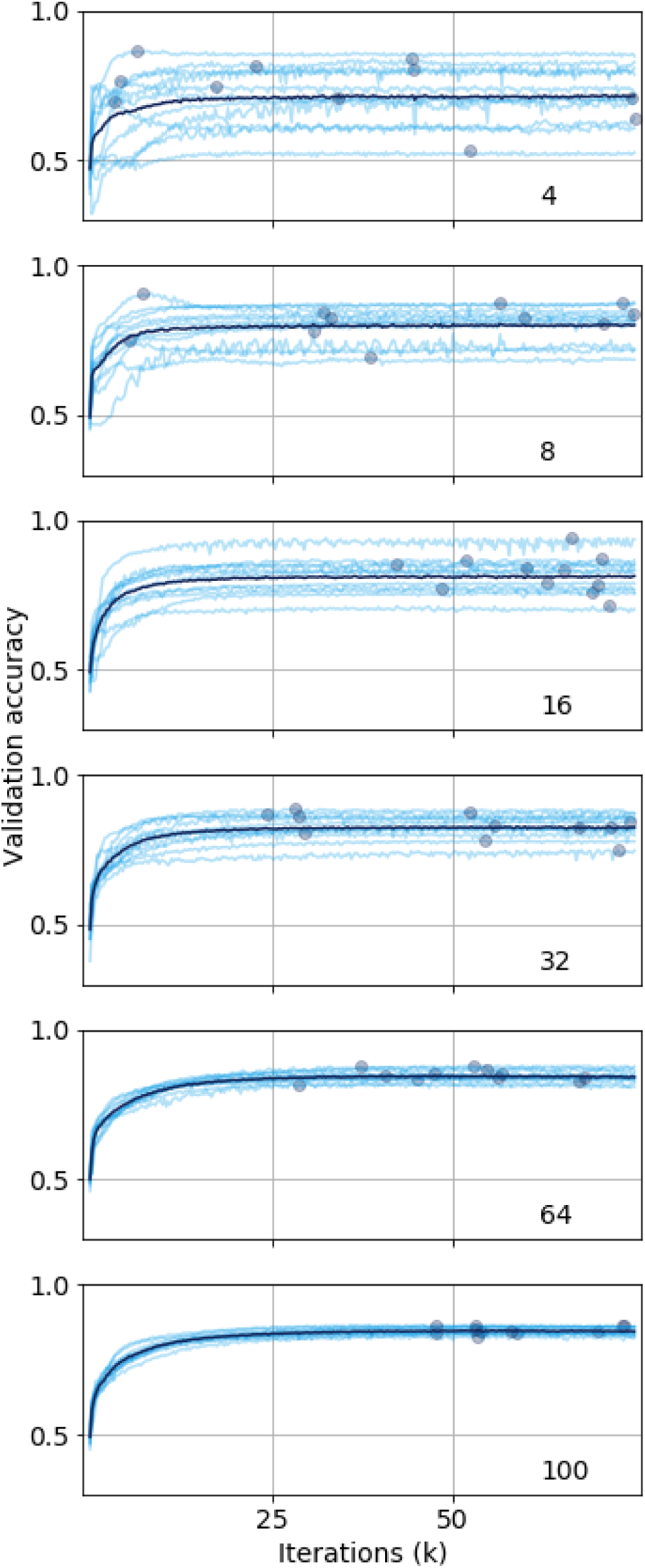
Learning curves for each subset size (percent). In each sub-plot, validation accuracy is plotted against number of training iterations. Each of the repetitions is plotted in light blue, and the average across repetitions is plotted in dark blue. Maximal validation accuracy in each course of training is plotted as a light blue point.

### C. Test accuracy as a function of subsample size

For each training run, the weights from the training history with the maximal validation accuracy were stored. The model was then assessed with these weights against the held out test set, yielding 11 test accuracy estimates for each proportion, as seen in Figure 2. As expected, test accuracy increased with sample size, reaching its maximal value at subsamples of 100 % (~86 % accuracy).

**Fig. 2.**
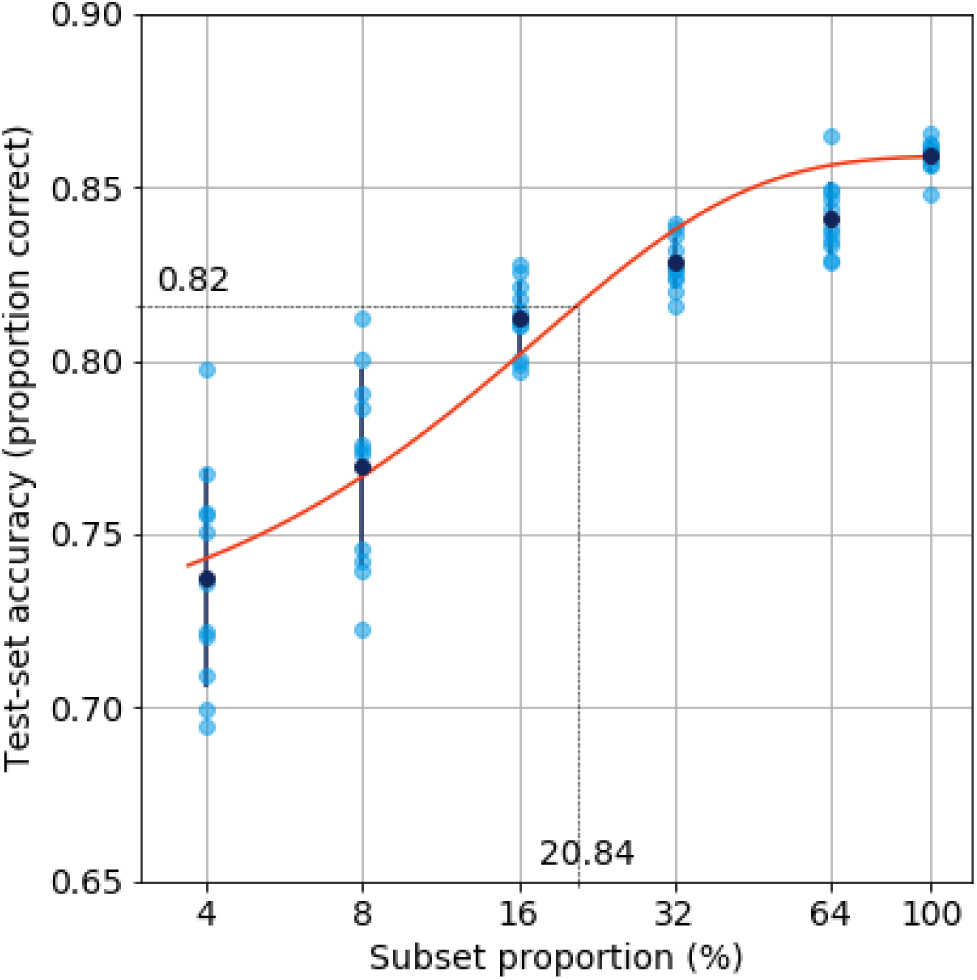
Accuracy on held out test set. Weights from the highest validation accuracy during training were used to test accuracy on a held out test set. Dark blue points are the means across 11 repetitions, with dark blue standard deviation error bars. Orange line: a logistic curve model was fit to all of the repetitions and sub-samples. According to this model, 95% of maximal accuracy (~82 % accuracy) can be achieved with 20.84% of the data (dashed lines).

Though accuracy is far from chance even with only 4 % of the data (~73% correct) it does increase precipitously between 4% and 64% of the data. However, it reaches close-to-maximal accuracy already at a proportion 16 - 32% of the total data-set. To quantify the amount of data needed to reach 95% of the maximal accuracy, we fit a two-parameter logistic function to the accuracy values across repetitions and sub-sample proportions (orange line in figure 2), fixing the saturation point of the function to be equal to the mean accuracy at subsamples of 100 % of the data. Inverting this function, we find that 95% of the maximal accuracy (approximately 82% accuracy) can already be achieved with 20% of the data (dashed lines in 2).

### D. What explains variability between repetitions?

Variability of test accuracy also diminished substantially with subsample size. Differences in variability in the comparisons across different proportions. These differences in variability could reflect two different factors: the first is the subsample size, and the other is the degree of overlap between different subsamples. For example, for 100 % subsamples, variability reflects only the random initial conditions of the network, because all subsamples of 100 % are identical. Similarly, the overlap between different subsets in higher proportions is likely to be larger than in smaller proportions. To evaluate the effect of this overlap, we conducted a separate experiment in which a single subset from 4% group was used, and training was repeated 11 times using the same set of images, to control for this effect. The learning curves from this protocol are are shown in Figure 3B, together with the learning curves from random subsamples (Figure 3B). The learning curves have similar variance as when random subsets are used and the maximal validation accuracy (Figure 3 C) does not differ between these protocols. This indicates that variability in test-set accuracy mostly relates to the size of the subsample, rather than to the amount of overlap between different subsets.

**Fig. 3.**
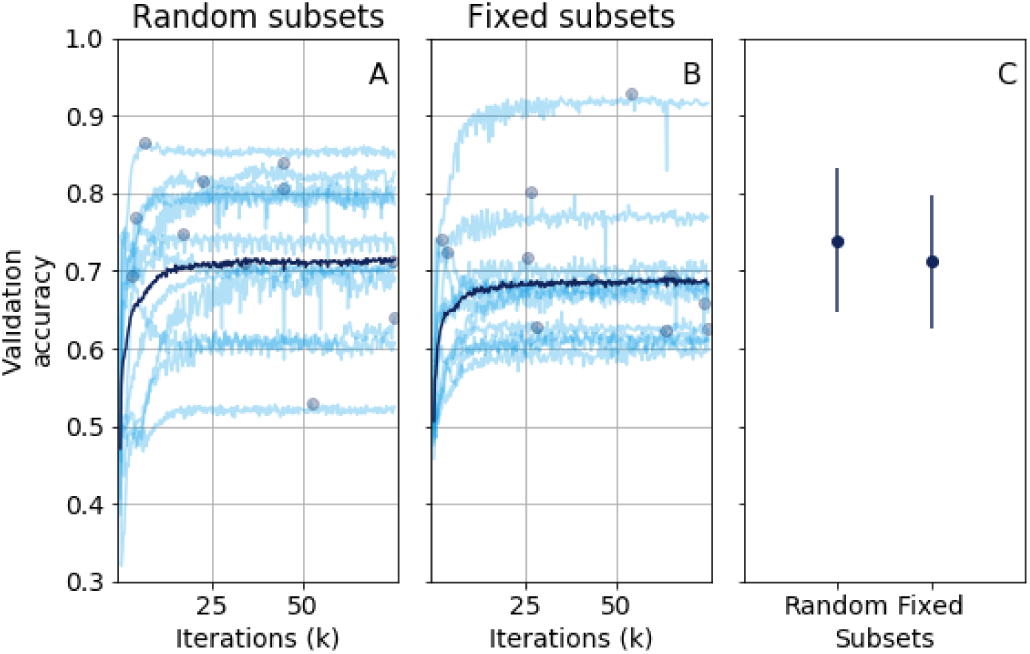
Learning curves of 4% subset. **A:** Random 4% subsets of the whole dataset chosen in each course of learning. **B:** Replications of the same 4% susample were repeated in each course of training. **C:** The average maximal validation accuracies (across the 11 courses of training) are plotted for random (left) and fixed (subsets) of 4% each, with standard deviation error bars

### E. The importance of a separate test set

To assess the importance of the separate test set, we computed the difference between the maximum validation accuracy and test set accuracy (Figure 4). We find that though variability in this difference decreases with larger sample size, there is no indication that this difference is larger for smaller sample sizes. In addition, there seems to be no overall bias indicating that the accuracy is systematically higher for the validation set, relative to the test set. This indicates that in the training protocol that we used, there was probably only minimal overfitting, or none at all.

**Fig. 4.**
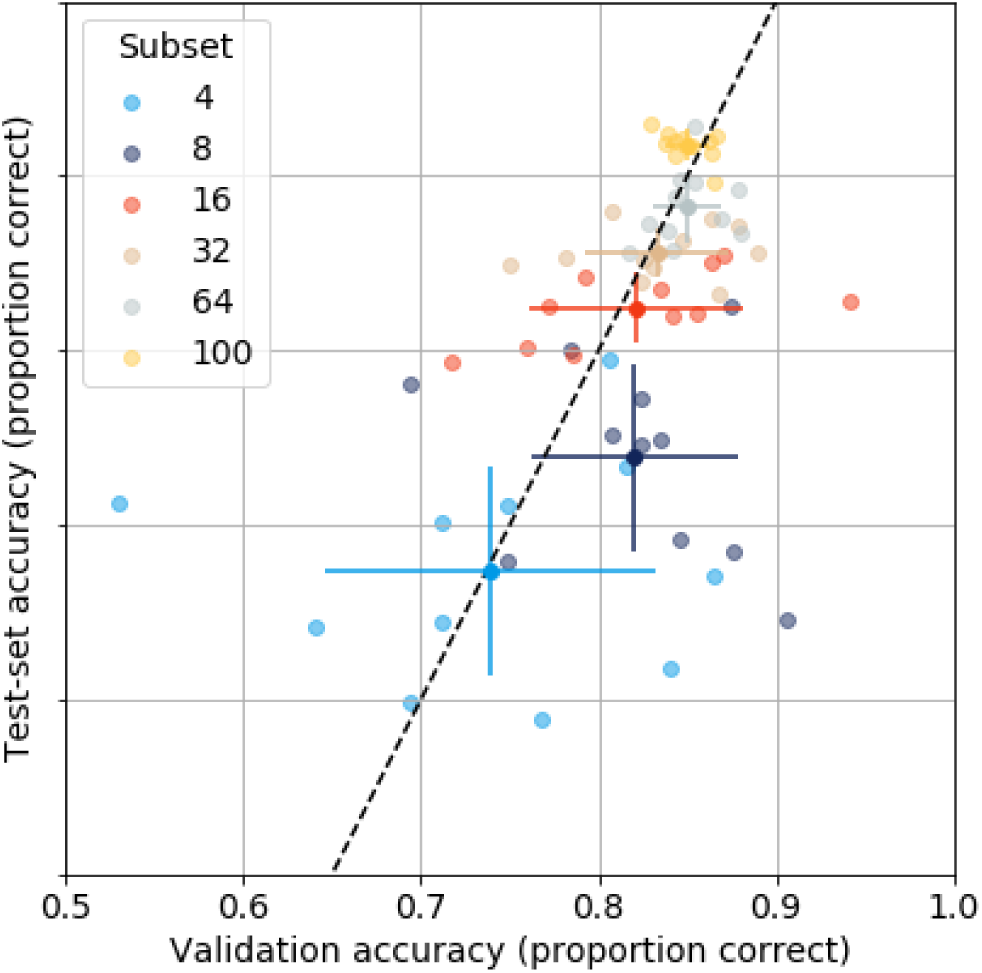
Difference between validation and test accuracy. Each light colored point represents a single course of training. The abscissa represents the maximal validation accuracy achieved during this course of training, while the ordinate represents the accuracy of the network in performing the classification on a separate test set, using the same weights that resulted in maximal validation accuracy. Solid colored points each represent the mean across 11 courses of training for each size subset, with standard deviation error bars. Dashed line indicates equality.

[fig3]

## IV. Conclusion

The application of DL in biomedical imaging is a promising avenue in current research. Future developments in this field may lead to accurate computer-assisted diagnosis systems with DL networks at their core. However, a major impediment to these developments is the assumption that DL requires very large data-sets that are not available for many types of nascent imaging technologies, or in the case of diseases that are relatively uncommon.

In the present work, we investigated the feasibility of DL with relatively small sample sizes. We focused on a prototype of computer-assisted diagnosis system that can accurately discriminate optical coherence tomography (OCT) images from retinae of patients with age-related macular degeneration AMD, relative to OCT images from the retinae of healthy controls.

### A. How many images do we need to discriminate AMD from healthy retina?

We found that training a DL network to perform at a high level of accuracy does not require millions of images. Instead, close-to-maximal performance is achieved with as few as approximately 20,000 images. Furthermore, the moderate increase in accuracy from 64% to 100% of the data suggests that further increase in data size would not result in much higher accuracy. This suggests that ~87% accuracy is as high as possible with these data and this DL algorithm. This limit on performance may be related to data quality; the partial accuracy of the labels in the EMR: these data are heterogeneous and ultimately depends on clinical decisions made by human observers, as well as the limited signal-to-noise ratio of the images in capturing the image features that are diagnostic. However, we do expect further performance improvements to come from more elaborate algorithms that incorporate additional information, or make better use of the information in the images, rather than only from more or better data.

The relatively small number of images required to train a DL network on this classification task is surprising given the VGG16 network has as many as 138M parameters [9]. Indeed, previous literature using similar networks (e.g. [2],[16]) used many millions of training samples to reach high accuracy. The discrepancy between our findings and the previous literature may stem from the differences between the use-case we present here, and the common use-cass for DL in previous literature. The assumption that many items from each class are required and that many millions of separate images are needed to train DL algorithms stems from the object classification literature mentioned in the introduction, but object classification in natural images addresses several challenges that are not typical in the classification of medical images, and particularly clinical images from OCT. Primary to these challenges is the variance in pose and orientation of natural objects within photographic images, which leads to large variance in the appearance of these objects. To capture all the variations of a category (e.g., ‘dog’), a DL network would have to be exposed to many thousands of exemplars of this category, generalizing not over all the angles from which this category could be captured, but also all the sub-categories of this category (e.g., ‘malamut’ or ‘poodle’). This variance in input is much more limited in biomedical images, such as OCT. In OCT images, the retina is always oriented in exactly the same direction, with the macula (the center of the retina) usually located in roughly the same part of the image. This reduces the complexity of training substantially, and we hypothesized that it might affect the data requirements for learning on data such as these.

Nevertheless, given the two-alternative classification task performed here, this finding is roughly consistent with the rule of thumb described by [17](“ … 5,000 labeled exemplars per category…”). The sub-sampling method introduced here provides a protocol for researchers that are interested in asking whether they have enough data to apply DL to their biomedical image data.

Note that because there are 11 images used in each OCT volume, the ~20,000 images represent approximately 1800 volumes of data, or 900 patients per group. This number of patients is well within range for many traditional random controlled trials and other clinical studies. This suggests that *de novo* training of DL networks could be integrated into many studies that are testing new imaging technologies, or that are studying less common disorders.

### B. Other strategies

There are currently two major alternative strategies to use for cases where data is limited. Data augmentation synthetically increases the sample size by performing transformations on the data [26]. This works well as long as the transformations performed to do not destroy the information necessary for classification, but introduce variability against which the DL network should develop tolerance. For clinical imaging data, examples of such transformations might be rigid translations and rotations of the image features.

The other strategy one might use when faced with limited data is transfer learning. This strategy is based on the observation that DL algorithms trained for different image processing tasks often learn very similar first-stage filters [27]. Therefore, in this approach learning begins with one (larger) data set. This dataset may share only some limited similarity to the datasets that are ultimately of interest, but this phase of learning allows the network to converge on good enough front-end filters. Once learning in this phase has converged, this network is then retrained on the dataset of interest. While this approach is promising, it is not clear what its limitations are, and whether it would work well for specific biomedical image processing tasks.

Both of these approaches are powerful complements to datasets that are not large enough, but even before employing these strategies, practitioners might want to assess whether the amount of data that they already have might be sufficient to accurately learn the classification task at hand.

### C. Do we need a separate test set?

Variance between different courses of training increased substantially with reduced sample sizes. This variance is not due to sampling of different individual items — both average accuracy and variance in accuracy between repetitions do not change substantially when the same items are repeatedly used in different subsamples.

Given the limit on test-set accuracy, one might expect that cross-validation on a separate test-set would be crucial, and that results in the test-set might differ substantially from the best performance on a validation set [28]. Nevertheless, we found no systematic difference in accuracy assessment on a separate test set relative to assessment of accuracy on a validation set that is repeatedly used during training. This finding is consistent with previous anecdotal evidence that has been mentioned in the literature [23].

Both of these findings may arise from the large number of parameters that are fit through the DL algorithm. The learning procedure may thus converge to different solutions based on initial conditions. When data is small, this may result in divergent solutions, that depend on the randomly generated initial conditions of the network. For larger networks, this implies that overfitting is not induced through repeated use of a validation data-set in accuracy assessments. However, further research would be needed to assess the limits of this conclusion and to merit its broad application.

## ACKNOWLEDGMENT

We would like to acknowledge NVIDIA Corporation for their generous donation of hardware in support of this research.A. Lee and Y. Wu were supported by an unrestricted grant by the Research to Prevent Blindness to the University of Washington. A. Rokem was supported through a grant from the Gordon & Betty Moore Foundation and the Alfred P. Sloan Foundation to the University of Washington eScience Institute Data Science Environment.

## Footnotes

1 “ As of 2016, a rough rule of thumb is that a supervised deep learning algorithm will generally achieve acceptable performance with around 5,000 labeled examples per category, and will match or exceed human performance when trained with a dataset containing at least 10 million labeled examples.” [17], page 20

